# Genomic characteristics, virulence and antimicrobial resistance in avian pathogenic *Escherichia coli* MTR_BAU02 strain isolated from layer farms in Bangladesh

**DOI:** 10.1101/2022.04.05.487091

**Authors:** Samina Ievy, M. Nazmul Hoque, Md. Saiful Islam, Md. Abdus Sobur, M. Shaminur Rahman, Fatimah Muhammad Ballah, Md. Bahanur Rahman, Jayedul Hassan, Mohammad Ferdousur Rahman Khan, Md. Tanvir Rahman

## Abstract

**Background:** Colibacillosis, caused by avian pathogenic *Escherichia coli* (APEC), is one of the most significant infectious diseases affecting poultry worldwide. APEC is one of the leading causes of mortality and morbidity associated with significant economic losses in the poultry industry.

**Objective:** This study was aimed to determine the genomic diversity, virulence factor genes (VFGs) and antimicrobial resistance (AMR) genes in the APEC isolated from layer chickens using whole-genome sequencing (WGS).

**Method:** APEC MTR_BAU02 strain was isolated from the sick and dead birds. Genomic DNA from APEC MTR_BAU02 strain was extracted using commercial DNA extraction kit, WGS libraries were prepared using the Nextera™ DNA Flex Library Prep Kit, and finally, paired-end (2 × 250) WGS performed using Illumina MiSeq sequencer.

**Results:** The genome size of strain APEC MTR_BAU02 is 4,924,680 bp with a GC content of 51.1% and 4,681 protein-coding sequences. Among the annotated WGS reads, 99.71% reads mapped to *Enterobacteriaceae* genomes. Based on the phylogenetic analysis of the APEC MTR_BAU02 genome and 99 reference genomes of *E. coli*, the APEC MTR_BAU02 genome showed sequence similarities with enterotoxigenic *E. coli* strains isolated from infections of different clinical severity. Metabolic functional annotations detected 380 SEED subsystems including genes coding for carbohydrate metabolism (17.34%), amino acid and derivatives (14.20%), protein metabolism (10.64%), cofactors, vitamins, prosthetic groups and pigments (7.49%), respiration (4.72%), membrane transport (4.49%), stress response (4.47%), motility and chemotaxis (4.46%), and virulence, disease and defense (2.22%). We also detected 92 VFGs and 122 AMR genes in the APEC MTR_BAU02 strain.

**Conclusion:** Assessment of these genomic features with functional genomic validation in commonly occurring serogroups of APEC will increase our understanding of the molecular pathogenesis, pave the way to early diagnosis and more effective control of colibacillosis, and improved animal welfare.

## 1. Introduction

*Escherichia coli* is a Gram-negative, dangerous, opportunistic pathogen in the poultry industry as it incurs significant economic losses from high mortality rates, reduced egg and meat production, and condemnations of whole flocks [1–4]. This ubiquitous pathogen has adaptive ability in diverse ecological niches, and can induce enteric and extraintestinal infections in animals and humans [3]. Moreover, strains of *E. coli* are thought to be virulent in humans and animals including commercial poultry and hold zoonotic potential [1, 5]. Colibacillosis, caused by *avian pathogenic E. coli* (APEC) is the most common bacterial diseases of chickens reared for egg and meat production [1, 3, 6], and poses a profound threat to one of humankind’s cheapest sources of high-quality animal protein [7]. This is one of the most significant infectious diseases affecting the global poultry industry [3, 8, 9]. APEC exists as a commensal component of the avian gut microbiota but emerges to take many forms, with a variety of systemic infections [3, 7]. Diseases of laying hens and other birds range from a syndrome associated with epidermal infections, mild to severe diarrhoea and/or enteritis, common respiratory tract (aerosacculitis) infections, perihepatitis, pericarditis, yolk sac infections, omphalitis and septicaemia [1, 3]. Though, different risk factors including chicken immunological immaturity and stress for colibacillosis have been identified, the disease has proved difficult to control. Moreover, strains of *E. coli* also have zoonotic significance since they are known to cause infections both in humans and animals including birds [3, 10]. Currently, one of the greatest challenges for APEC is the difficulty of understanding their pathogenicity [4]. In addition, the pathogenicity appears to be linked to a heterogenous mix of plasmid and chromosomal genes involved in bacterial adhesion, invasion, toxicity, antibiotic resistance, survival and metabolism under stress [1, 11], and the emergence of virulent APEC from a background of commensal gut-dwelling *E. coli* [1, 12].

Antimicrobial resistance (AMR) is an ever-increasing public health crisis [13–15]. The G20 partners have recognized AMR as a major “growing threat to public health and economic growth” [4]. It causes an estimated 700,000 deaths each year across the world. Drug-resistant APEC strains can contaminate the food supply from farm to fork through eggs, meat, and other contaminants and thus pose a severe threat to the consumer’s health [3, 4]. The indiscriminate use of antibiotics in poultry production may have contributed to drug resistance in APEC. From the poultry farm, drug-resistant strains are deposited into soil, wastewater, air, and the environment. The relation of antibiotics resistance especially in APEC phylotypes had previously been discussed [3, 16]. There are reports that resistant *E. coli* strains might enhance antimicrobial resistance (AMR) in other organisms (pathogenic and non-pathogenic) within gastrointestinal tract of the chicken [3, 16], and help in transmitting and disseminating multidrug resistant (MDR) strains, and antimicrobial resistance (AMR) genes between animal and human pathogens [3]. Many of the earlier studies have explored the association between pathogenic traits of APEC, and their virulence factors associated genes (VFGs) repertoire [40–43]; however, potential pathogenic traits associated with APEC strains warrant the attention into the potential role of VGFs, MDR properties, and their cross-talk with specific host. Technological advent in high-throughput whole-genome sequencing has dramatically revolutionize the clinical microbiology to investigate the population genomics of pathogen evolution [1]. However, the understanding the spread of APEC in avian population remains challenging for two reasons such as uncertainty about the transmission of a few globally distributed epidemic clones [17] or a diverse assemblage of disease-causing lineages [11], and the genes contributing to APEC virulence are less well described than in human pathotypes [1].

A series of studies from many countries have detected drug resistance determinants in APEC [3, 4, 18, 19], however, most of these studies did not focus on APEC and the associated virulence genes. In order to investigate the role of APEC in the pathophysiology of poultry as well as its contributions in spreading AMR genes and VFGs, the reported avian pathogenic *E. coli* MTR_BAU02 (APEC MTR_BAU02) strain was isolated from different poultry farm of Bangladesh. Here, we report the whole genome of the APEC MTR_BAU02 strain, and its functional genomics to elucidate the mechanisms through which this pathogen shows its resistant and virulent phenotypes, contributes to AMR genes and VFGs spreading and pathogenesis in layer farms in Bangladesh.

## 2. Materials and Methods

### 2.1. Ethics statement

The experimental procedures and protocols used in this study were approved by the Animal Welfare and Experimentation Ethics Committee of Bangladesh Agricultural University (approval number AWEEC/BAU/2019(28)).

### 2.2. Sample collection and processing

A total of 99 samples including air from the insides of poultry shades (n = 31), feces from sick birds (n = 32), and the internal organs (trachea, intestine, liver, lung, and egg yolk material; n = 36) of dead birds were collected aseptically from 32-layer farms located in Mymensingh district, Bangladesh during January–November 2019. The settle plate method was used for air sampling following previously described by protocols [20] with some modifications. In brief, instead of nutrient agar, here, eosin methylene blue (EMB) agar plates were exposed at 1 m above the ground to different corners of the poultry shades for 10 min. We collected the freshly dropped fecal samples using sterile cotton buds from sick isolated groups of birds. Internal organs were collected during post-mortem examinations. All the collected samples were given unique tag numbers and transported to the laboratory maintaining the cold chain. Immediately after arrival at the laboratory, fecal samples (1 g) were seeded into test tubes containing 5 mL of nutrient broth. Internal organs were initially cut into small pieces and then transferred into test tubes containing 5 mL of nutrient broth. EMB agar plates and test tubes were then incubated aerobically at 37 °C overnight.

### 2.3. Isolation and identification of *E. coli*

The isolation and identification of *E. coli* was based on culture on EMB agar plates following previously published protocols [3, 4]. In brief, one loopful inoculum from previously prepared nutrient broth (NB) containing each sample was inoculated into freshly prepared NB composed of yeast extract-2 gm/l, peptone-5 gm/l, and sodium chloride-5 gm/l, and incubated for 24 h at 37°C. A small amount of inoculum from NB was streaked onto EMB agar for 24 h at 37°C for selective growth. The colonies showing characteristic metallic sheens on the EMB agar plates (3–5 colonies from each sample) were considered as indicative of *E. coli*. These colonies were then subjected to morphological study by Gram staining, biochemical tests (indole, methyl-red, catalase, citrate and Voges–Proskauer) for confirmatory identification of *E. coli*. The final confirmation of the isolation of *E. coli* was performed by polymerase chain reactions (PCRs) targeting the *E. coli* 16S rRNA gene [4].

### 2.4 Genomic DNA extraction

The genomic DNA from APEC MTR_BAU02 strain was extracted using commercial DNA extraction kit, QIAamp DNA Mini Kit (QIAGEN, Hilden, Germany) according to the manufacturer’s instruction. The DNA concentration and purity was measured by a NanoDrop 2000 UV-Vis Spectrophotometer (Thermo Fisher, Waltham, MA, USA). DNA extracts with A260/280 and A260/230 ratios of ~ 1.80 and 2.00 to 2.20, respectively, were considered as high-purity DNA sample. Finally, the harvested DNA was visualized on 1% (w/v) agarose gel, and chosen for DNA sequencing based on their high purity and adequate concentration.

### 2.5. Library preparation and whole genome sequencing

DNA libraries were prepared using the Nextera™ DNA Flex Library Prep Kit (Illumina, San Diego, CA, USA) according to the manufacturer’s protocol. The *E. coli* library profile showed similarities with that of the Reference Guide of the Nextera™ DNA Flex Library Prep Kit. The library quality was also strengthened by the electrophoresis results that showed each library having a fragment size of approximately 300–1000 bp. Thus, it was demonstrated that the quality of the *E. coli* libraries met the Illumina (Illumina, San Diego, CA, USA) platform requirements. Whole genome sequencing (WGS) of the prepared libraries was performed through Illumina MiSeq sequencer (Illumina, San Diego, CA, USA) using a 2 × 250 paired-end protocol.

### 2.6 Genome assembly and data analysis

The FASTQ reads quality was initially assessed by the FastQC tool [21] followed by trimming of low-quality residues at read beginnings and endings (‘leading:10’ and ‘trailing:10’), Nextera XT adaptors, other Illumina specific sequences (‘Illuminaclip’ set to value ‘NexteraPE-PE.fa:2:30:10’), and reads less than 200 bp using the Trimmomatic [22], while the quality cut off value was Phred-20 [23]. Trimmed reads were de novo assembled using SPAdes 3.15.2 [24], setting the ‘–careful’ and ‘–cov-cutoff auto’ options to reduce mismatches and short indels, and remove low coverage contigs, respectively. Orphaned reads resulting from trimming (i.e., reads where only one read of the pair survived) were provided to the assembler as single-end reads. Subsequently, trimmed paired-end and orphaned reads were mapped against the de novo assembly, the Sakai *E. coli* O157:H7 reference genome (NCBI accession NC_002695.2) [25] using Bowtie2 2.3.0 [26] with the ‘–sensitive’ and ‘–end-to–end’ settings. KmerFinder 3.1 [27] and PathogenFinder 1.1 [28] opensource tools were utilized to identify the species (i.e., *E. coli*) and pathogenicity of the isolates (MTR_BAU02).

### 2.7 Comparative genome analysis

APEC MTR_BAU02 genome was compared with the reference strains APECO1 (GenBank accession number NC_008563) [29], and APECO78 (GenBank accession number CP004009) [30] genomes. APEC isolates were obtained from lesions of chickens and turkeys clinically diagnosed with colibacillosis from various locations within the United States [29, 30]. The circular visualization of the APEC MTR_BAU02 genome was performed using CGViewer [31]. We further used the NCBI Tree Viewer (https://www.ncbi.nlm.nih.gov/tools/treeviewer/) and MEGA v7.0 [32] to perform the whole genome-based phylogenetic analysis applying maximum-likelihood method with 1000 bootstraps. The phylogenetic tree was subsequently imported to iTOL (v. 3.5.4) (http://itol.embl.de/) [33] for better visualization, and bootstrap values were reported for each branch.

### 2.8 Genomic functional potentials analysis

Draft genome of APEC MTR_BAU02 strain was initially annotated using RAST (Rapid Annotation using Subsystem Technology) server [34], and KEGG (Kyoto Encyclopedia of Genes and Genomes) Automatic Annotation Server (KAAS) [35]. The RAST server provided data on the distribution of genes in various categories [23]. The ResFinder [36, 37], and VirulenceFinder [36, 37] databases were used to detect AMR genes and virulence genes, respectively. We utilized SnapGene Viewer web tool (https://www.snapgene.com/snapgene-viewer/) to visualize the virulence plasmid of the APEC MTR_BAU02. The ResFinder database integrated within AMR++ pipeline identified the respective genes or protein families coding for the AMR gene markers in the APEC MTR_BAU02 strain [23].

### 2.9 Data availability

The whole genome shotgun (WGS) of the APEC MTR_BAU02 strain has been deposited at GenBank under accession number JAIWJB000000000, and the assembly reports of the genome are also available at GenBank. The version described in this paper is version JAIWJB000000000.1. The Illumina shotgun reads are available in the National Center for Biotechnology Information (NCBI) Sequence Read Archive (SRA) accession number SRR16193628 under BioProject accession number PRJNA766218.

## 3. Results

In this study, 82.83% (82/99) were positive for *E. coli* according to the selective EMB culture and PCR targeting of the *E. coli* 16S rRNA gene from clinically diagnosed colibacillosis cases of laying chickens. The whole genome sequence (WGS) analysis of the study isolate, MTR_BAU02 using KmerFinder 3.1 detected the isolate as *E. coli*, while the pathogenicity of the isolate was confirmed (0.951 out of 1.00, close to the pick value indicating higher pathogenicity) by PathogenFinder 1.1.

### 3.1 Genome features of the APEC MTR_BAU02 strain

The de novo genome assembly revealed that the APEC MTR_BAU02 genome was 4,924,680 bp in size with 4,497,218 bp DNA coding sequences, and a coverage of 38.98x. The draft genome had 2,516,511 bp DNA sequences assigning for G+C content of 51.1% (Table 1). The complete genome of APEC MTR_BAU02 strain possessed 94 contigs including 332,120 bp contig N50, and 6 contig L50. Of the detected contigs, the largest and smallest contig size were 4,49,968 bp and 234 bp, respectively (Fig. S1). Notably, preliminary sequence analysis revealed 41 insertion sequences (ISs), 14 predicated genomic islands (GIs), seven prophage-related sequences, two CRISPR arrays, and one plasmid (Table 1). The genome of APEC MTR_BAU02 encodes for 4,801 genes including 4,681 protein coding sequences (CDSs), 28 rRNAs (5S = 8, 16S = 5, 23S = 15), 81 tRNAs, 11 ncRNAs, and 88 pseudogene (Fig. 1). The presumptive SEED functions of the 70 putative genes were predicted by comparing with the non-redundant protein sequences in NCBI database. The physical map of these genes detected in APEC MTR_BAU02 genome is illustrated in Fig. 2. Fifty-one of the 70 predicted genes (72.86%) showed high sequence homology to known functional genes with identified SEED functions, ten genes (14.29%) denoted for extracytoplasmic sensor or ligand binding domain (LBD) having potential roles in signal transduction, while eight genes (11.43%) encoded for conserved hypothetical proteins with unknown functions and one gene (1.43%) coded for putative autotransporter (Fig. 2).

**Table 1.** Avian pathogenic *E. coli* (APEC) strain MTR_BAU02 genome characteristics.

| Attribute(s) | Value | % of Total<sup>a</sup> |
| --- | --- | --- |
| Genome size (bp) | 4,924,680 | 100.00 |
| DNA coding (bp) | 4,497,218 | 91.32 |
| Genome coverage | 38.98x | 38.98 |
| DNA G + C (bp) | 2,516,511 | 51.1 |
| Total contigs | 94 |  |
| Contig N50 (bp) | 332,120 | 6.74 |
| L50 | 6 |  |
| Total genes | 4,803 | 100.00 |
| Coding sequences (CDSs) | 4,681 | 97.46 |
| Protein coding genes | 4,571 | 95.17 |
| RNA genes | 120 | 2.5 |
| tRNA genes | 81 | 1.69 |
| rRNA genes | 28 | 0.58 |
| Pseudo genes | 112 | 2.33 |
| Genes with function prediction | 3,940 | 82.03 |
| Genes assigned to SEED subsystems | 3,512 | 73.12 |
| Number of subsystems | 380 |  |
| Genes with Pfam domains | 4,214 | 87.74 |
| Genes with signal peptides | 386 | 8.04 |
| Genes with transmembrane helices | 1,084 | 22.57 |
| CRISPR arrays | 2 |  |
| Number of plasmid | 1 |  |

<sup>a</sup>Total based on either the size of the genome in base pairs (bp) or the total number of genes in the annotated genome.

**Fig. 1.**
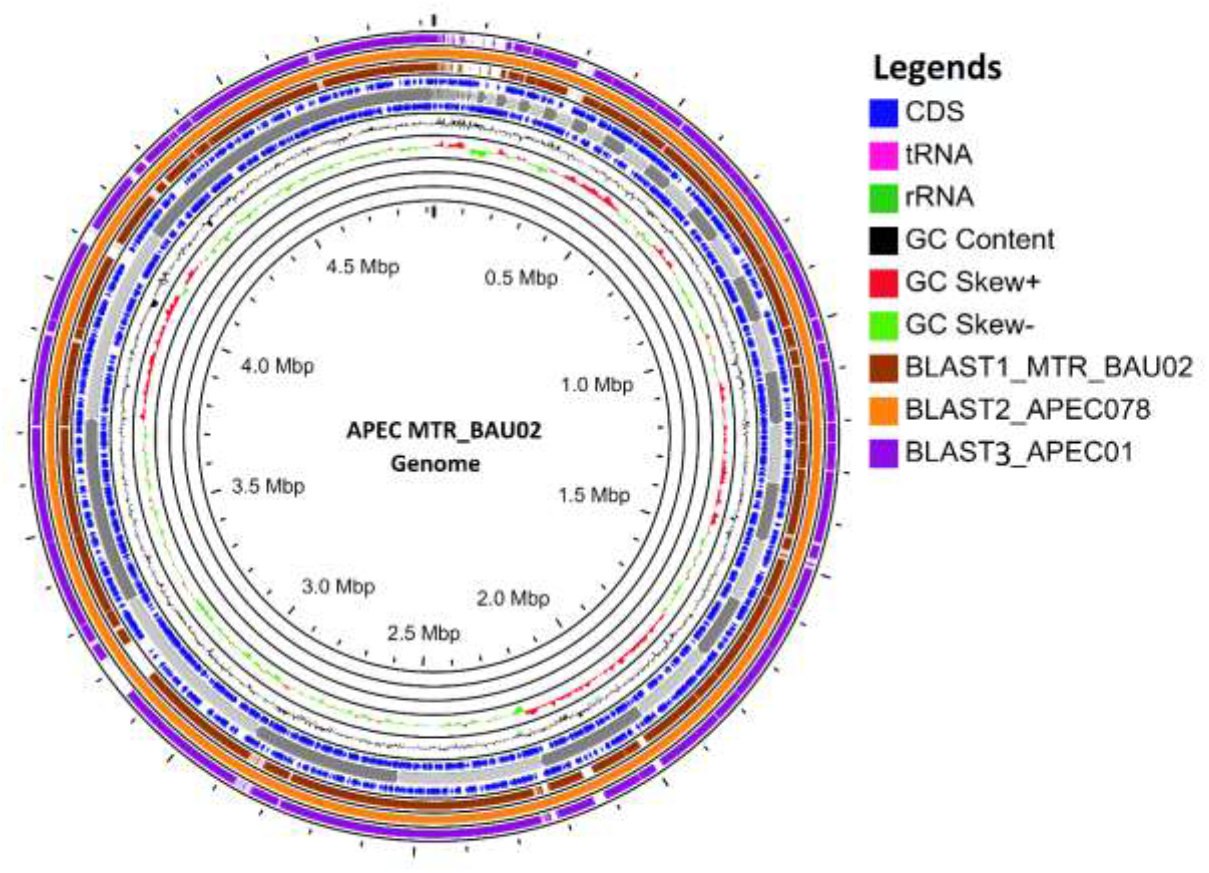
Circular genome representation of APEC MTR_BAU02 compared with APEC01 (NC_008563) and APEC078 (CP004009). The outermost ring: *E. coli* blast 3 results for APEC01(purple) followed by *E. coli* blast 2 results APEC078 (solid yellow), *E. coli* blast 3 results MTR_BAU02 (purple) representing the positions covered by the BLAST comparative alignment results (BLASTN), G+C content (Black), G+C positive skew (red), and G+C negative skew (green). Image created using CGview Server.

**Fig. 2.**
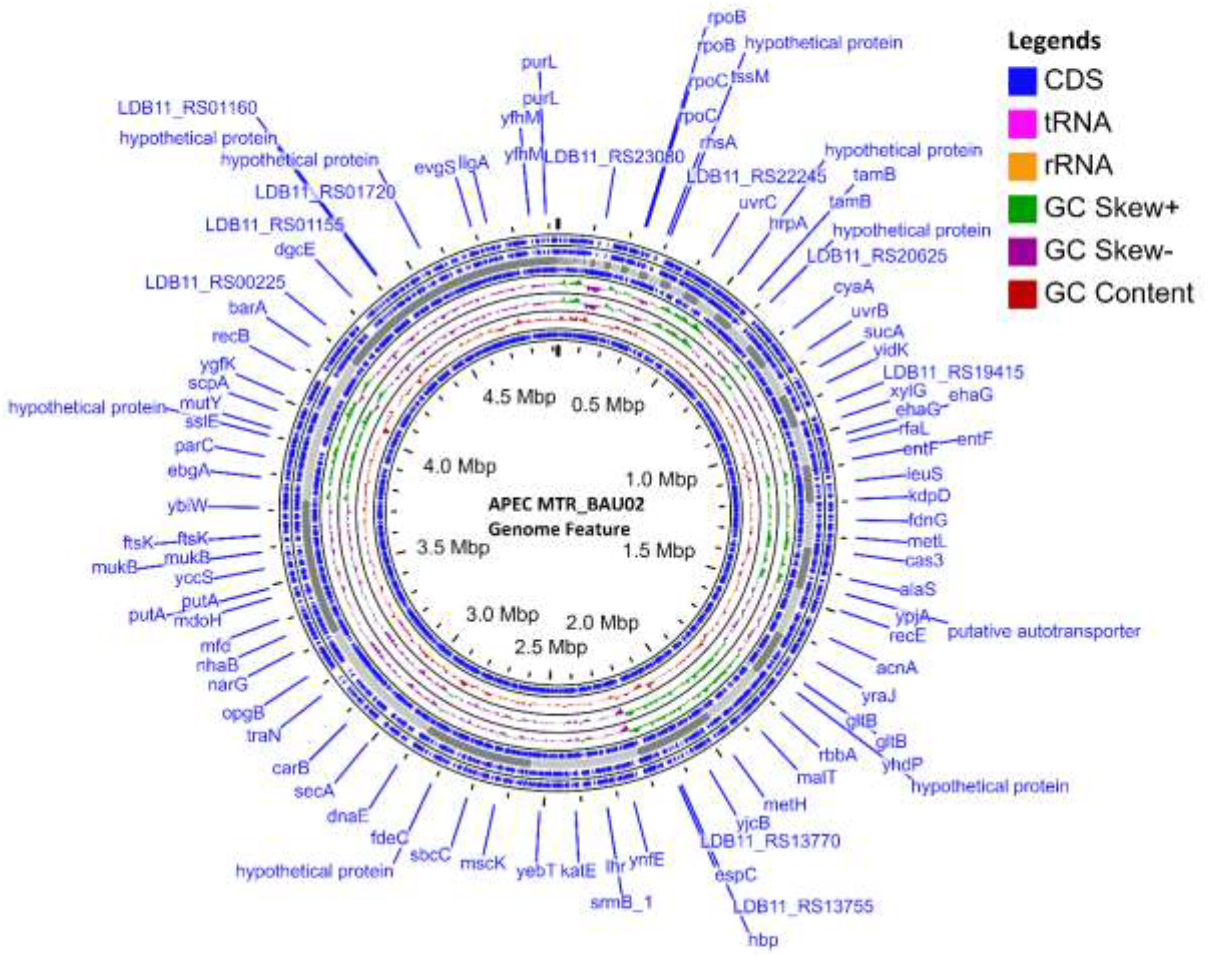
BRIG- and CGView-derived schematic map of circular genome of the APEC MTR_BAU02. The ORFs with predicted annotations are indicated with blue arrows in the innermost ring which shows the genome location. The innermost ring is followed by rings representing GC content (dark red), GC skew + (green) and — (purple), respectively. The two most external rings show identified coding sequences (CDS) on both strands are represented in blue arrows and results of genome annotation process. The putative function of each CDS is labelled at the outermost ring and the organization of the SEED functional modules of 70 in the CDS are represented at the innermost ring.

### 3.2 Phylogenetic relatedness of the APEC MTR_BAU02 strain

To interpret the evolution of APEC MTR_BAU02 strain, a selection of 34 *E. coli* complete genomes downloaded from NCBI was used to map phylogenetic tree by using maximum-likelihood method. All complete genomes of *E. coli* except our APEC MTR_BAU02 genome were named as the NCBI uid. The results showed that the *E. coli* genomes were separated into three major clades (I, II, and III). Clades I and II were further subdivided into four subclades. APEC MTR_BAU02 genome showed the closest evolutionary relationship with APEC O1 (Fig. 3), indicating that APEC MTR_BAU02 may have evolved from avian samples of colibacillosis.

**Fig. 3.**
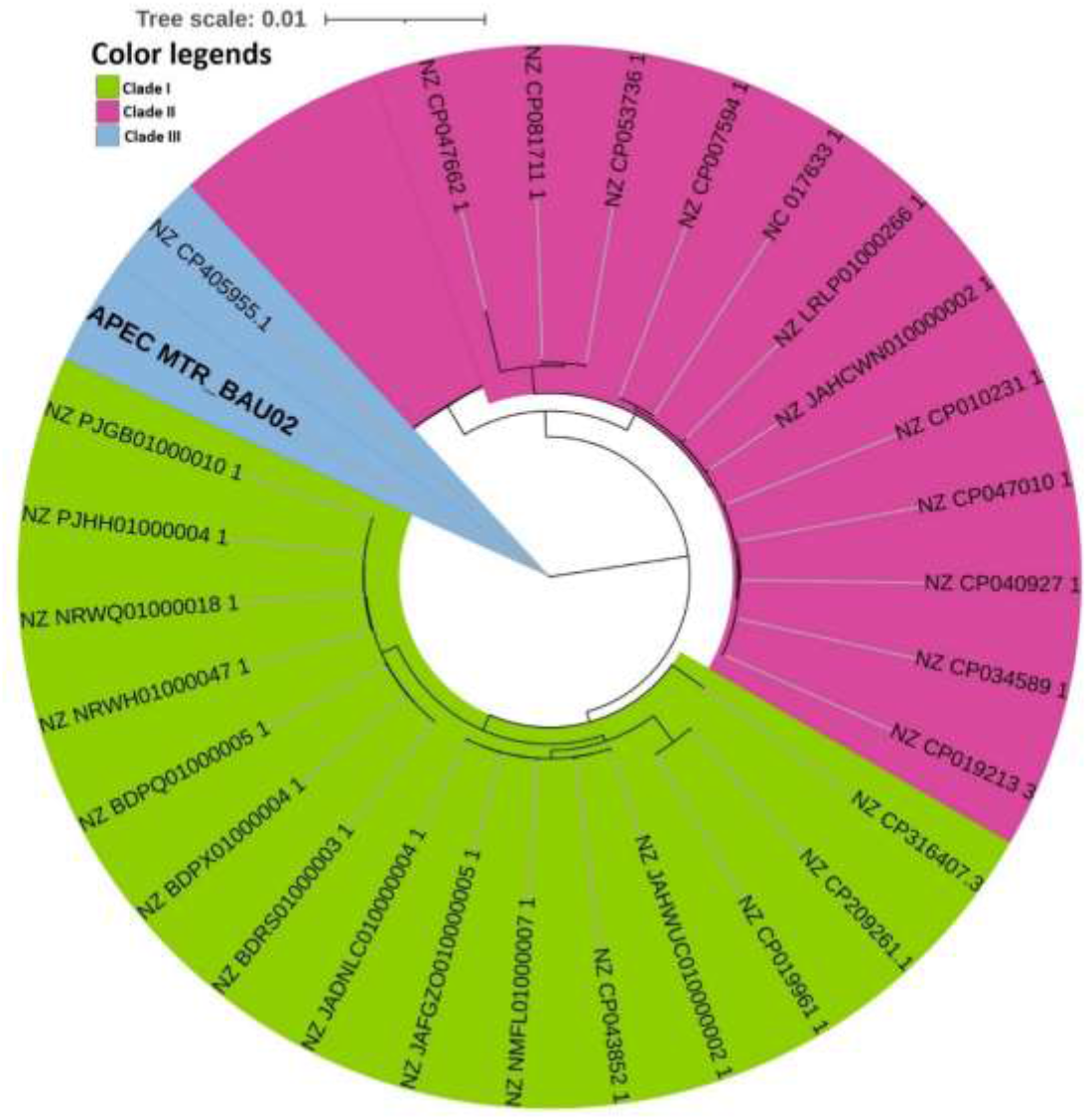
The Evolutionary relationships between APEC MTR_BAU02 and other *E. coli* strains. Thirty-four strains of *E. coli* with complete genomes from NCBI were used for phylogenetic analysis. The mid-point rooted tree was constructed using the NCBI Tree Viewer (https://www.ncbi.nlm.nih.gov/tools/treeviewer/), and visualized with iTOL (interactive tree of life). The evolutionary relationship was inferred using the maximum-likelihood method. Different colors are assigned according to the close evolutionary relatedness (clade) of the genomes. The scale is in the unit of the number of substitutions per site.

Among the annotated WGS reads, 99.71% reads mapped to *Enterobacteriaceae* genomes (i.e., *E. coli*). Based on the phylogenetic analysis of the APEC MTR_BAU02 genome and 99 *E. coli* reference genomes, the APEC MTR_BAU02 genome showed highest sequence similarities with enterotoxigenic *E. coli* (MP020980.1) strain (4.0%) followed by *E. coli* (2846750, 2780750, p0305293.5 etc) strains (3.0% identity with each genome) (Fig. 4) isolated from infections of different clinical severity.

**Fig. 4.**
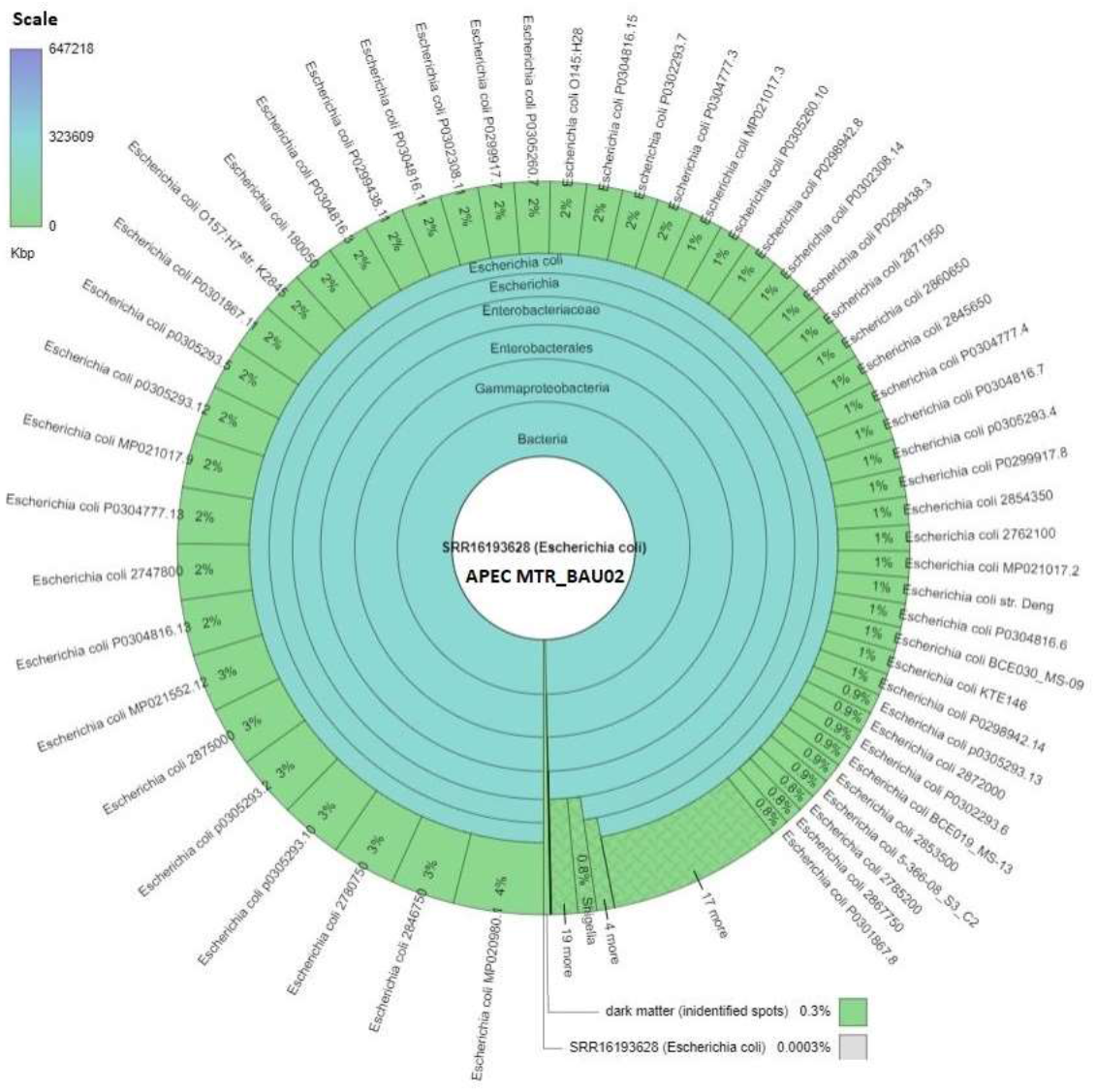
A snapshot of a Krona hierarchical representation of 100 *E. coli* complete genomes along with APEC MTR_BAU02 genome. A multi-layered interactive version with zoom and six-step depth adjustment show the taxonomic hierarchical similarities of APEC MTR_BAU02 genome with rest of the *E. coli* genomes retrieved from NCBI. The APEC MTR_BAU02 genome (SRR16193628) genome showed more than 99% sequence similarities with other *E. coli* genomes. The scale bar represents the length of *E. coli* genomes (Kbp).

### 3.3 Metabolic functional features in the APEC MTR_BAU02 genome

The detected functional pathways were distributed under 380 SEED subsystems, which were mostly represented by the genes coding for carbohydrate metabolism (CHO; 17.34%), amino acid and derivatives (14.20%), protein metabolism (10.64%), cofactors, vitamins, prosthetic groups and pigments (7.49%), respiration (4.72%), membrane transport (4.49%), stress response (4.47%), motility and chemotaxis (4.46%), and virulence, disease and defense (2.22%). Rest of the SEED subsystems had relatively lower abundances (< 2.0%) (Fig. 5). Investigating deeper into these SEED subsystem distribution, we found that genes coding for monosaccharide (23.2%) and central CHO (23.18%) metabolism, protein biosynthesis (68.26%), and lysine, threonine, methionine, and cysteine metabolism (30.46%) were top abundant metabolic functional categories in the APEC MTR_BAU02. Moreover, flagellar motility (100%), oxidative stress (38.14%), type II protein secretion system (23.71%), chaperon pathway (16.50%), electron donating reactions (55.88%), folate and pterions metabolism (29.63%), fatty acids, lipids, and isoprenoids (54.54%) were the predominating metabolic features in APEC MTR_BAU02 (Fig. 5).

**Fig. 5.**
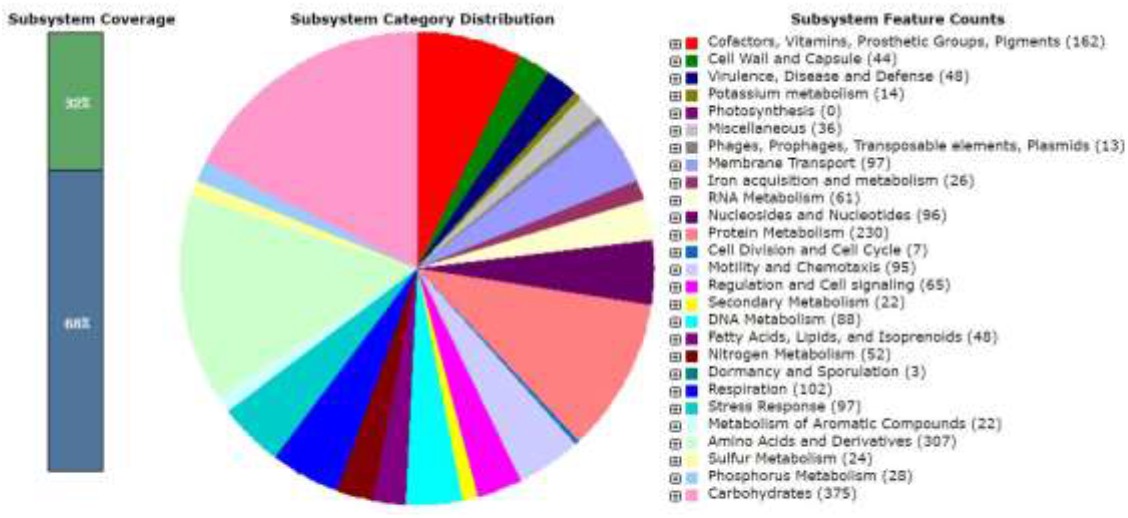
Subsystem categorical distribution in APEC MTR_BAU02 genome. The whole genome sequence of the APEC strain MTR_BAU02 was annotated using the Rapid Annotation System Technology (RAST) server. The pie chart demonstrates the count of each subsystem feature and the subsystem coverage.

### 3.4 Virulence and antimicrobial resistance genes

The APEC MTR_BAU02 genome encoded different genes, proteins and/or pathways coding for virulence. The APEC MTR_BAU02 genome possessed a virulence plasmid of 73,864 bp (Fig. 6). The virulence plasmid of APEC MTR_BAU02 strain possessed a 627 bp region that comprises characteristics of flagellar assembly protein (fliH), 591 bp region of imidazole glycerol phosphate synthase subunit (hisH), 261 bp region of Rz1-like lysis system protein (lysC), 210 bp region for heme exporter protein (ccmD), 834 bp region for sulfate/thiosulfate ABC transporter permease (cysT), and 321 bp region coding for bifunctional 3-phenylpropionate/cinnamic acid dioxygenase ferredoxin subunit (hcaC) (Fig. 6). Using VFDB database, 92 VFGs were detected in the APEC MTR_BAU02 genome. Among the identified VFGs, genes coding for iron uptake (25.58%), type II secretion system (20.47%), invasion (16.41%), adherence (12.83%), motility (10.33%), fimbrial activity (7.45%), serum resistance (2.17%), autotransporter (1.80%), and quorum sensing system (1.80%) were the most predominant virulence classes in the APEC MTR_BAU02 genome. Rest of the VFGs had relatively lower (<1.0%) abundances (Fig. 7).

**Fig. 6.**
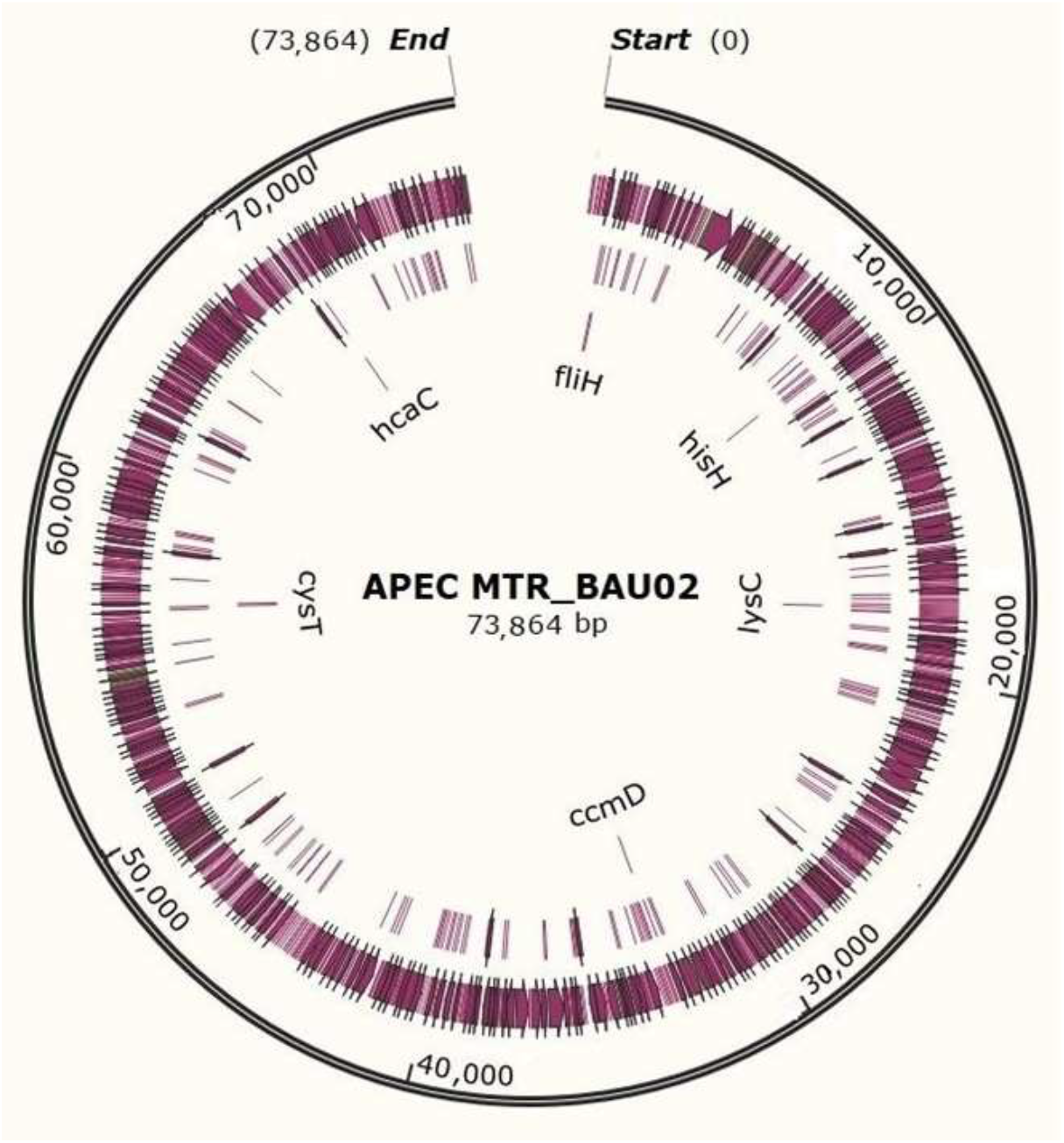
Map of the virulence plasmid of the APEC MTR_BAU02. The outermost two rings indicate the size of the plasmid. The next circle represents all putative open reading frames (ORFs), depending on ORF orientation clockwise and anticlockwise directions, respectively. The inner most ring shows predicted virulence features extracted from the genome GenBank file of APEC MTR_BAU02. The map was created by using SnapGene Viewer.

**Fig. 7.**
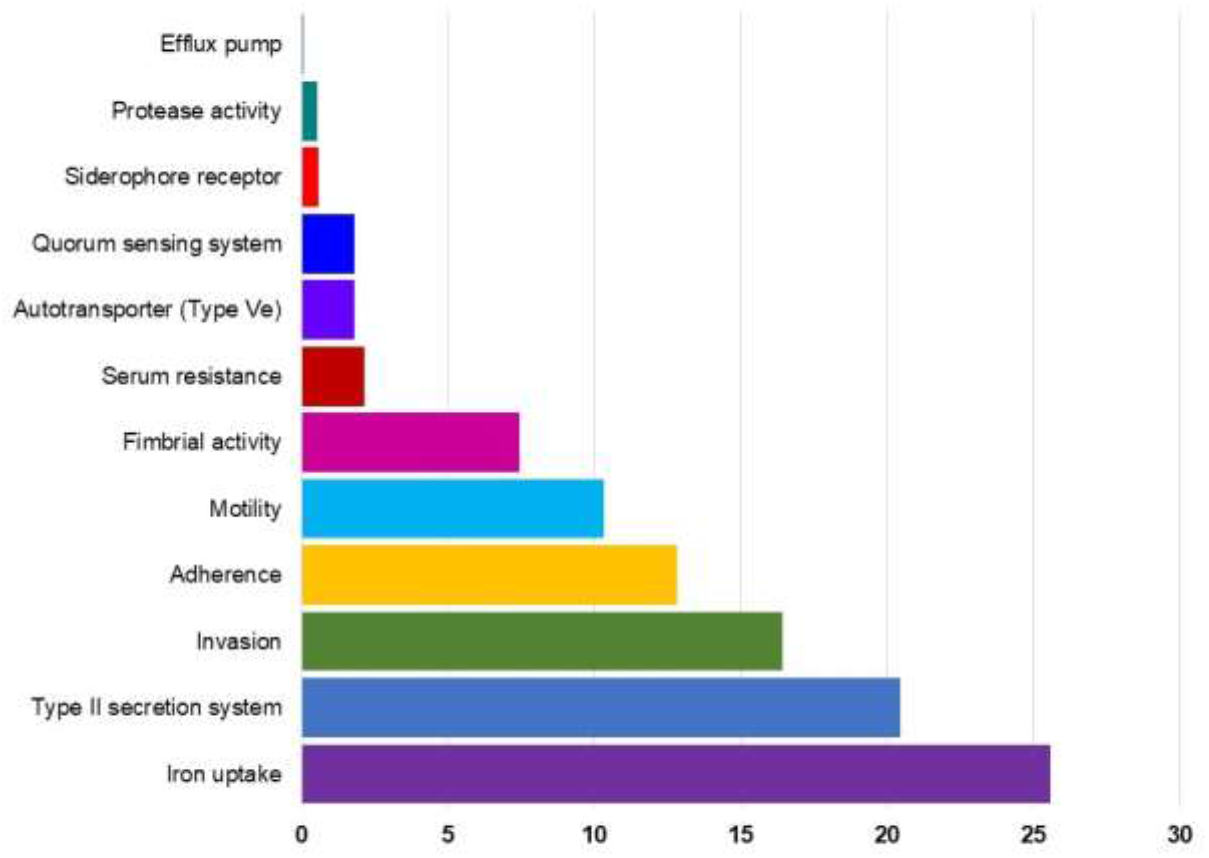
An overview of the virulence factor genes (VFGs) assigned to the genome of APEC MTR_BAU02. The complete genome sequence of the APEC MTR_BAU02 strain was annotated using the VirulenceFinder database. The bar plots demonstrate the count of each VFG feature and the coverage.

We also detected 122 genes coding for antimicrobial resistance (AMR) using the ResFinder database (Fig. 8). The top abundant genes were *Mdt*ABC-*Tol*C (7.77%), *Evg*A (3.88%), *Emr*AB-OMF (3.88%), *Emr*KY-*Tol*C (3.88%), *Acr*D (2.91%), *Acr*AD-*Tol*C (2.91%), *Acr*EF-*Tol*C (2.91%), *Mdt*C (2.91%), *Emr*B (2.91%) and *MdtB* (2.91%). Rest of the genes had relatively lower abundances (< 2.0%) (Fig. 8). These genes were related to several important genomic functions of APEC MTR_BAU02 conferring resistance to multiple antibiotics, and of them, multi-drug efflux pump system, transcriptional activator, multi-drug efflux pump system_membrane fusion component, probable aminoglycoside efflux pump, intrinsic mechanism of multidrug resistance, multidrug resistance efflux pump outer membrane protein, and multidrug resistance proteins. Therefore, efflux pump conferring resistance to multiple antibiotics was found as the predominating resistance class in the genome of APEC MTR_BAU02.

**Fig. 8.**
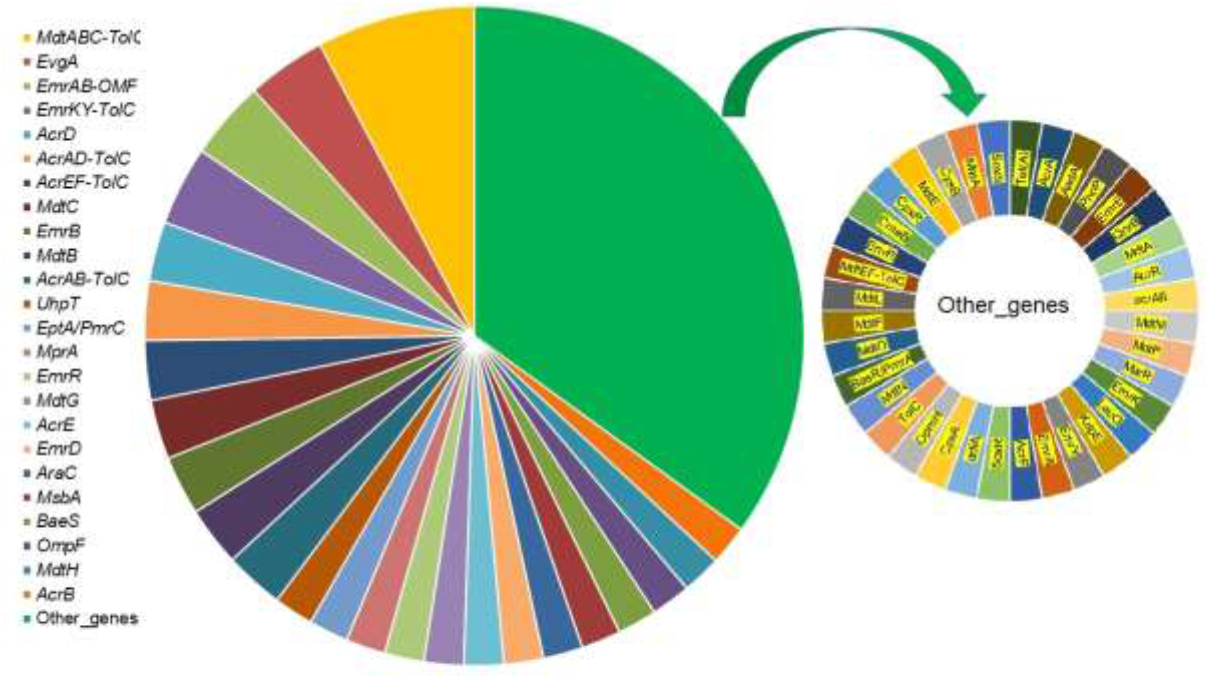
Antimicrobial resistance (AMR) genes assigned to the genome of APEC MTR_BAU02. The complete genome sequence of the APEC MTR_BAU02 strain was annotated using the ResFinder database. The Venn diagram demonstrate the count of each AMR feature and the coverage.

## 4. Discussion

*E. coli* are simultaneously ubiquitous in animal guts and a major cause of diverse intestinal and extraintestinal infections [1]. The avian colibacillosis is an acute and globally occurring infectious disease of domestic and wild birds caused by different strains of pathogenic *E. coli*, and it is associated with considerable economic losses mainly due to the morbidity and mortality associated [3]. Therefore, isolation and identification *E. coli* from both healthy chicken guts and infected systemic tissues (APEC) allowed characterisation of the genotypes and gene pools associated with different sites, and health states. APEC-associated avian colibacillosis has a significant impact on the poultry industry, and globally, poultry industries are being confronted by enormous economic losses due to the dramatic impact of the disease [3, 4, 23]. APEC are also important from the public health point of view. To control colibacillosis, multiple antimicrobials have been used indiscriminately, especially in middle- and low-income countries including Bangladesh, contributing to the development and spread of AMR [14, 23]. The subsequent selection of MDR strains has generated serious challenges in terms of public health. Sustainable development goals (SDGs) are affected by AMR, especially in targeting hunger, poverty, malnutrition, health, and economic growth [14, 38]. Thus, investigations of APEC strains with regard to their phylogenetic evolution, metabolic functional potentials, virulence genes (VFGs) and AMR profiles may help to curtail their hazardous effects.

The complete genome of APEC strain MTR_BAU02 was determined by initial *de novo* assembly. The features of the complete genome of APEC MTR_BAU02 described in this study is consistent with the genome composition of different APEC strains reported earlier in several studies [1, 39, 40]. Phylogenetic analysis using complete genomes (WGS) provides a greater resolution that helps to determine the evolutionary relatedness among isolates. This type of analysis allows automated and more robust epidemiological surveillance [41, 42]. In this study, phylogenetic analysis of the 34 *E. coli* genomes was performed to comparing the evolutionary relatedness of the strains. The result of phylogeny showed that 34 *E. coli* strains could be clearly divided into three monophyletic clades, which was similar to the whole-genome-based phylogeny of many of the previous studies [23, 30, 39, 43]. However, it is important to note that prokaryotic genomes are highly influenced by the reference genome selected as well as the closeness between isolates being analysed. In this regard, we selected the genomes of APECO1 (GenBank accession number NC_008563) [29], and APECO78 (GenBank accession number CP004009) [30] as a reference genomes for comparing strains. The APEC MTR_BAU02 strain showed its closest evolutionary relationship with another APEC genome (APEC01 and APEC078) reported from USA [1]. In this study, APEC MTR_BAU02 was the most divergent genome compared to its closest available complete genomes retrieved from the database. The conclusion to be drawn from the lack of correlations is that firstly APEC are very diverse and secondly it is not possible to rely on any one or more of the assembly method and tests for reconstructing phylogenetic tree to define APEC. In our analysis, genomes were grouped based on their allelic profile and serotype. In our study, we defined cluster complexes among genomes of similar origin, but this not necessarily means that those isolates were closely related.

Plasmids have been shown to play an important role in the pathogenicity of most *Enterobacteria*. In this study, we detected a virulence plasmid in the APE MTR_BAU02 genome. Plasmids occur commonly among APEC strains, and may be a defining feature of the APEC pathotype [44]. Despite the fact that this plasmid contains many of the genes or proteins thought to contribute to *E. coli* virulence, their acquisition by APEC MTR_BAU02 strain may not necessarily result in increased virulence of the laying birds. These finding also corroborated with the findings of several earlier studies [44–46]. We also detected 92 VFGs in the APEC MTR_BAU02 isolate, including genes coding for iron uptake, type II secretion system, bacterial invasion, adherence, motility, fimbrial activity, serum resistance, autotransporter, and quorum sensing system. These VFGs are crucial in the pathogenesis of various types of infections caused by different pathotypes of *E. coli* [23, 30, 47].

Apart from the wide genetic diversity, we detected a high prevalence of the AMR genes in the APEC MTR_BAU02 isloate. We identified 122 AMR genes, of which *Mdt*ABC-*Tol*C, *Evg*A, *Emr*AB-OMF, and *Emr*KY-*Tol*C showed the highest prevalence. Overexpression of mdtABCD (and separately of mdtABC) in *E. coli* confers increased resistance to deoxycholate, cholate, taurocholate, novobiocin, nalidixic acid, norfloxacin, fosfomycin, and other antibiotics and toxic compounds [48]. The response regulator *Evg*A controls expression of multiple genes conferring antibiotic resistance in *E. coli* [49]. *Emr*AB is the first multidrug efflux transporter identified in *E. coli*, constitutively expressed in *E. coli* and contributes to intrinsic resistance against various antimicrobial compounds [50]. Moreover, using WGS in this study provided more information on resistance to multiple antibiotics, through multi-drug efflux pump system, transcriptional activator, multi-drug efflux pump system_membrane fusion component, probable aminoglycoside efflux pump, intrinsic mechanism of multidrug resistance, multidrug resistance efflux pump outer membrane protein, and multidrug resistance proteins. This concordance is consistent with other studies and also supports the available evidence on the robustness of WGS in predicting AMR phenotypes [51]. Interestingly, MDR efflux genes and proteins were detected in APEC MTR_BAU02 isolate in this study. Although, specific efflux systems for specific antimicrobial classes such as aminoglycosides, bicyclomycin, and macrolides were not observed in this study, other efflux systems, especially the MdtABC-TolC, AcrAB-TolC, and AcrAD-TolC system, were observed in the APEC MTR_BAU02 isolate and have been reported to confer MDR to tetracycline, chloramphenicol, ampicillin, nalidixic acid, and rifampicin [47, 51]. These finding the prevalence AMR genes in APEC MTR_BAU02 genome suggests that the commensal *E. coli* population in poultry in countries with a high and unregulated use of antibiotics constitutes a hotspot for selection for carriage of transferrable resistance. This is worrisome and calls for immediate action to reduce the preventive use of antimicrobials in general and colistin in particular. One of the major health issues associated with *E. coli* is its role in the emergence and the dissemination of antimicrobial resistance [52]. Colibacillosis in the poultry farms might be prevented and/or controlled by the rational therapeutic use of antimicrobials. However, evolution of MDR APEC strains along with the transmission of resistance genes has created challenges in reducing the risk of APEC infections [3]. This study has some limitations. First, the study was conducted using only the sequence based WGS data which did not allow us to observe the identified VFGs and AMR genes through laboratory culture and PCR amplification. Second, the study was conducted in a certain geographical region in Bangladesh. The addition of more farms from different regions of the country would bring a more detailed picture of the *E. coli* diversity. Third, only layer chickens were investigated in this study, whereas broiler and backyard chickens are also a major part of poultry production in Bangladesh. Finally, although the study identified diverse VFGs and AMR genes in *E. coli* from layer birds, however, it was not possible to clarify whether the source of the virulent and resistant-strains was a vertical transmission from breeder flocks or not.

## Conclusion

This is the first WGS based study on multidrug-resistance APEC in layer chicken in Bangladesh. The WGS provided in this study will also allow for comparative genomic analysis with other *E. coli* isolates from different sources, which may help to elucidate the dissemination mechanisms of AMR genes among bacteria, animals, and humans. These results call for immediate action from the policy makers to stop imprudent use or to actively regulate the rational use of antibiotics along with other critically important antimicrobials in poultry production in Bangladesh.

---

**Fig. S1.**
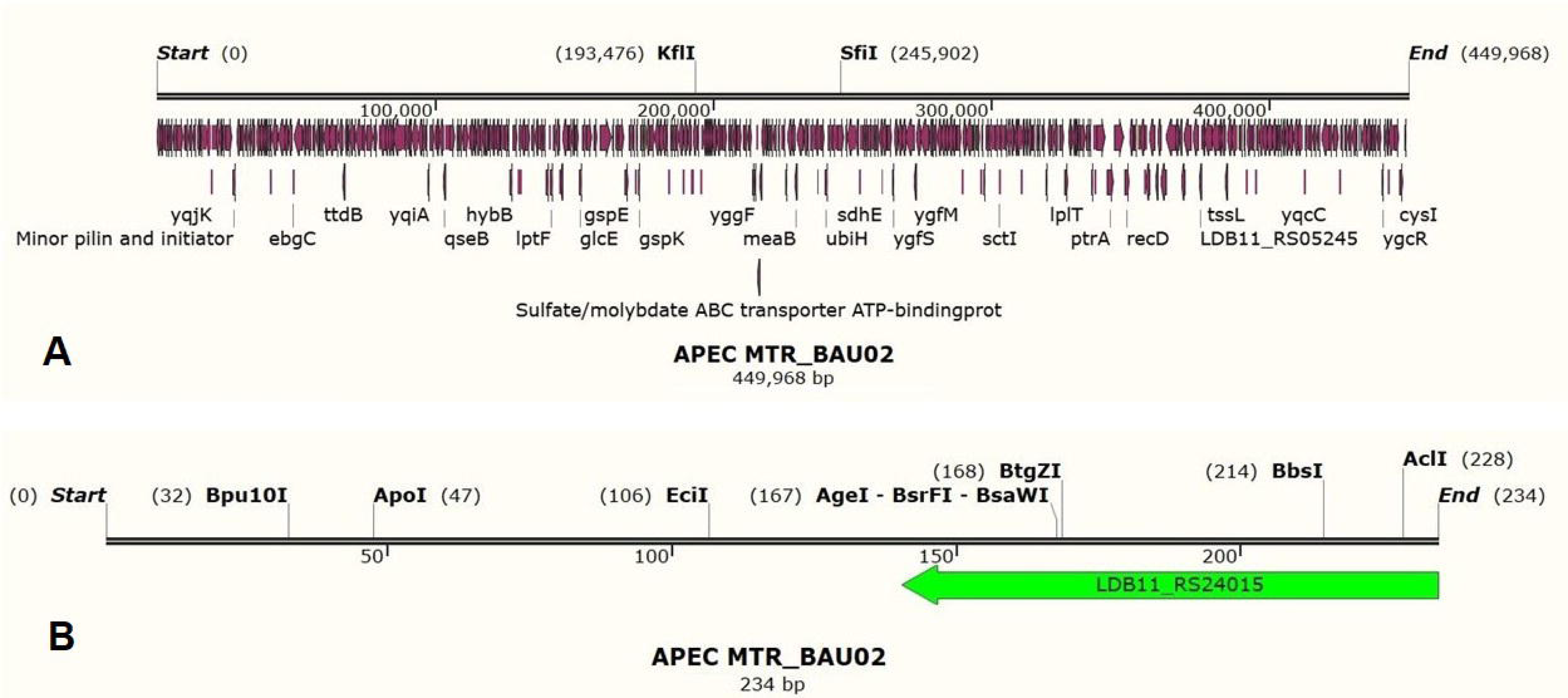
Contigs of the APEC MTR_BAU02. A. The largest contig (4,49,968 bp) in the WGS of the APEC MTR_BAU02; B. The smallest contig (234 bp) in the APEC MTR_BAU02 genome.

## Acknowledgments

We are deeply grateful to the Ministry of Education, Government of Bangladesh for funding the research project (Project No. LS2018686). In addition, we are grateful to the poultry farm owners for giving access to the samples. The authors would also like to thank the Department of Microbiology and Hygiene, Faculty of Veterinary Science Bangladesh Agricultural University, Mymensingh for the support during the research.

## Ethical approval

Not required.

